# A*PA2: up to 20 times faster exact global alignment

**DOI:** 10.1101/2024.03.24.586481

**Authors:** Ragnar Groot Koerkamp

## Abstract

**Methods:** We introduce A*PA2, an exact global pairwise aligner with respect to edit distance. The goal of A*PA2 is to unify the near-linear runtime of A*PA on similar sequences with the efficiency of dynamic programming (DP) based methods. Like Edlib, A*PA2 uses Ukkonen’s band doubling in combination with Myers’ bitpacking. A*PA2 1) extends this with SIMD (single instruction, multiple data), 2) uses large block sizes inspired by Block Aligner, 3) avoids recomputation of states where possible as suggested before by Fickett, 4) introduces a new optimistic technique for traceback based on diagonal transition, and 5) applies the heuristics developed in A*PA and improves them using *pre-pruning*.

**Results:** The average runtime of A*PA2 is 19*×* faster than the exact aligners BiWFA and Edlib on *>*500 kbp long ONT reads of a human genome having 6% divergence on average. On shorter ONT reads of 11% average divergence the speedup is 5.6*×* (avg. length 11 kbp) and 0.81*×* (avg. length 800 bp). On all tested datasets, A*PA2 is competitive with or faster than approximate methods.

**Availability:** github.com/RagnarGrootKoerkamp/astar-pairwise-aligner

## 1 Introduction

The problem of *global pairwise alignment* is to find the shortest sequence of edit operations (insertions, deletions, substitutions) to convert a string into a second string (Needleman and Wunsch, 1970; Vintsyuk, 1968), where the number of such operations is called the *Levenshtein distance* or *edit distance* (Levenshtein, 1965; Wagner and Fischer, 1974).

Over time, the length of genomic reads has increased from hundreds of basepairs to hundreds of thousands basepairs now. Meanwhile, the complexity of practical exact algorithms has not been improved by more than a constant factor since the introduction of the diagonal transition algorithm (Ukkonen, 1985; Myers, 1986).

Our recent work A*PA (Groot Koerkamp and Ivanov, 2024) uses the A* shortest path algorithm to speed up alignment and has near-linear runtime when divergence is low. A drawback of A* is that it uses a queue and must store all computed distances, causing large (up to 500×) overhead compared to methods based on dynamic programming (DP).

This work introduces A*PA2, a method that unifies the heuristics and near-linear runtime of A*PA with the efficiency of DP based methods. As Fickett (1984, p. 1) stated 40 years ago and still true today, at present one must choose between an algorithm which gives the best alignment but is expensive, and an algorithm which is fast but may not give the best alignment.

In this paper we narrow this gap and show that A*PA2 is nearly as fast as approximate methods.

### 1.1 Contributions

We introduce A*PA2, an exact global pairwise sequence aligner with respect to edit distance.

In A*PA2, we combine multiple existing techniques and introduce a number of new ideas to obtain up to 19× speedup over existing single-threaded exact aligners. Furthermore, A*PA2 is often faster and never much slower than approximate methods.

As a starting point, we take the band doubling algorithm implemented by Edlib (Šošić and Šikić, 2017) using bitpacking (Myers, 1999). First, we speed up the implementation (points 1., 2., 3.). Then, we reduce the amount of work done (4., 5.). Lastly, we apply A* heuristics (6.).

**1. Block-based computation**. Edlib (Fig. 1d) computes one column of the DP matrix at a time, and for each column decides which range (subset of rows) of states to compute. We significantly reduce this overhead by processing blocks of 256 columns at a time (Fig. 1e), taking inspiration from BLOCK ALIGNER (Liu and Steinegger, 2023). Correspondingly, we only store states of the DP-matrix at block boundaries, reducing memory usage.

**Fig. 1.**
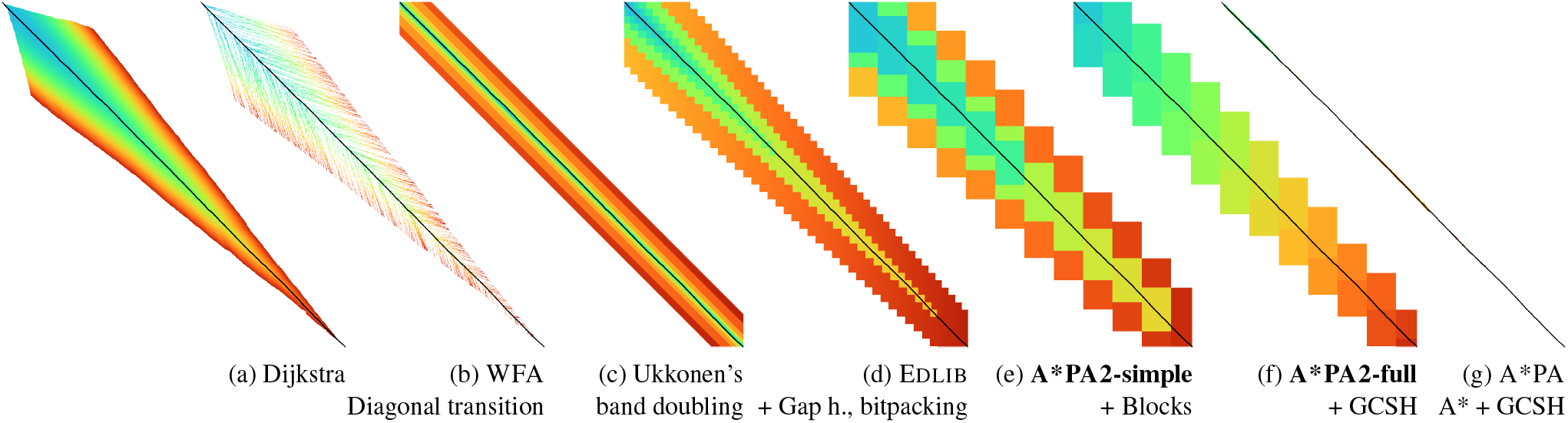
Alignment of two sequences of length 3000 bp with 20% divergence using different algorithms and methods. Coloured pixels correspond to visited states in the edit graph or dynamic programming matrix, and the blue to red gradient indicates the order of computation. The black path indicates the optimal alignment. Gap h. stands for the gap heuristic and GCSH for the gap-chaining seed heuristic of A*PA. (c) to (f) incrementally add features.

**2. SIMD**. We speed up the computation of each block by using 256bit SIMD, allowing the processing of 4 computer words in parallel.

**3. Novel encoding**. We introduce a novel encoding of the input sequence to speed up SIMD operations by comparing characters bit-by-bit and avoiding slow gather instructions. This limits the current implementation to alphabets of size 4.

**4. Incremental doubling**. Both the band doubling method of Ukkonen (1985) and Edlib recompute states after doubling the threshold. We avoid this by using the theory behind the A* algorithm, extending the incremental doubling of Fickett (1984) to blocks and arbitrary heuristics.

**5. Traceback**. For the traceback, we optimistically use the diagonal transition method (Ukkonen, 1985; Myers, 1986; Marco-Sola *et al*., 2020) within each block with a strong adaptive heuristic, only falling back to a full recomputation of the block when needed.

**6. A\***. We apply the gap-chaining seed heuristic (GCSH) of A*PA (Figs. 1f and 1g) (Groot Koerkamp and Ivanov, 2024), and improve it using *pre-pruning*. This technique discards most of the *spurious* (off-path) matches ahead of time.

### 1.2 Previous work

We give a brief overview of developments that this work builds on, in chronological order per approach. See also, e.g., the reviews by Kruskal (1983) and Navarro (2001), and the introduction of our previous paper Groot Koerkamp and Ivanov (2024). Section 2 covers relevant topics more formally.

#### Needleman-Wunsch

Pairwise alignment has classically been approached as a dynamic programming problem. For input strings of lengths *n* and *m*, this method creates a (*n* + 1) × (*m* + 1) table that is filled cell by cell using a recursive formula. Needleman and Wunsch (1970) gave the first *O*(*n*^2^*m*) algorithm, and Sellers (1974) and Wagner and Fischer (1974) improved this to what is now known as the *O*(*nm*) *Needleman-Wunsch algorithm*, building on the quadratic algorithm for *longest common subsequence* by Sankoff (1972).

#### Graph algorithms

It was already realized early on that an optimal alignment corresponds to a shortest path in the *edit graph* (Vintsyuk, 1968; Ukkonen, 1985). Both Ukkonen and Myers (1986) remarked that this can be solved using Dijkstra’s algorithm (Dijkstra, 1959), taking *O*(*ns*) time (Fig. 1a), where *s* is the edit distance between the two strings and is typically much smaller than the string length. (Although Ukkonen only gave a bound of *O*(*nm* log(*nm*)).) However, Myers (1986, p. 2) observes that the resulting algorithm involves a relatively complex discrete priority queue and this queue may contain as many as *O*(*ns*) entries even in the case where just the length of the […] shortest edit script is being computed.

Hadlock (1988) realized that Dijkstra’s algorithm can be improved upon by using A* (Hart *et al*., 1968), a more *informed* algorithm that uses a *heuristic* function *h* that gives a lower bound on the remaining edit distance between two suffixes. He uses two heuristics, the widely used *gap cost* heuristic (Ukkonen, 1985; Hadlock, 1988; Wu *et al*., 1990; Spouge, 1989, 1991; Papamichail and Papamichail, 2009) that simply uses the difference between the lengths of the suffixes as lower bound (Fig. 1d), and a new improved heuristic based on character frequencies in the two suffixes. A*PA (Groot Koerkamp and Ivanov, 2024) improves the *seed heuristic* of Ivanov *et al*. (2021) to the *gap-chaining seed heuristic* (GCSH) with *pruning* to obtain near-linear runtime when errors are uniform random (Fig. 1g). Nevertheless, as Spouge (1991, p. 3) states, algorithms exploiting the lattice structure of an alignment graph are usually faster and further (Spouge, 1989, p. 4):

This suggests a radical approach to A* search complexities: dispense with the lists [of open states] if there is a natural order for vertex expansion.

In this work we follow this advice and replace the plain A* search in A*PA with a much more efficient approach based on *computational volumes* that merges DP and A*.

#### Computational volumes

Wilbur and Lipman (1983) is, to our knowledge, the first paper that speeds up the *O*(*nm*) DP algorithm, by only considering states near diagonals with many *k-mer matches*, but at the cost of giving up the exactness of the method. Fickett (1984) notes that for some chosen parameter *t* that is at least the edit distance *s*, only those DP-states with cost at most *t* need to computed. This only requires *O*(*nt*) time, which is fast when *t* is an accurate bound on the distance *s*. For example *t* can be set as a known upper bound for the data being aligned, or as the length of a suboptimal alignment. When *t* = *t*_0_ turns out too small, a larger new bound *t*_1_ can be chosen, and only states with distance in between *t*_0_ and *t*_1_ have to be computed. When *t* increases by 1 in each iteration, this closely mirrors Dijkstra’s algorithm.

Ukkonen (1985) introduces a very similar idea, statically bounding the computation to only those states that can be contained in a path of length at most *t* from the start to the end of the graph (Fig. 1c). On top of this, Ukkonen introduces *band doubling*: *t*_0_ = 1 can be doubled (*t*_*i*_ = 2^*i*^) until *t*_*k*_ is at least the actual distance *s*. This find the alignment in *O*(*ns*) time.

Spouge (1989) unifies the methods of Fickett and Ukkonen in *computational volumes* (see Section 2): small subgraphs of the full edit graph that are guaranteed to contain all shortest paths. As Spouge notes:

The order of computation (row major, column major or antidiagonal) is just a minor detail in most algorithms.

But this is exactly what was investigated a lot in the search for more efficient implementations.

#### Parallelism

In the 1990s, the focus shifted from reducing the number of computed states to computing states faster through advancements in implementation and hardware. This resulted in a plethora of new methods. While there many recent methods optimizing the computation of arbitrary scoring schemes and affine costs (Smith and Waterman, 1981; Gotoh, 1982; Bergeron and Hamel, 2002; Suzuki and Kasahara, 2018; Shao and Ruan, 2024), here we focus on methods for computing edit distance.

The first technique in this direction is *microparallelism* (Alpern *et al*., 1995), also called SWAR (SIMD within a register), where each (64 bit) computer word is divided into multiple (e.g. 16 bit) parts, and word-size operations modify all (4) parts in parallel. This was then applied with *inter-sequence parallelism* to align a given query to multiple reference sequences in parallel (Alpern *et al*., 1995; Baeza-Yates and Gonnet, 1992; Wu and Manber, 1992; Hyyrö *et al*., 2005; Rognes, 2011). Hughey (1996) notes that *anti-diagonals* of the DP matrix are independent and can be computed in parallel, to speed up single alignments. Wozniak (1997) applied SIMD (single instruction, multiple data) for this purpose, which are special CPU instructions that operate on multiple computer words at a time. Rognes and Seeberg (2000, p. 702) also use microparallelism, but use *vertical* instead of anti-diagonal vectors:

The advantage of this approach is the much-simplified and faster loading of the vector of substitution scores from memory. The disadvantage is that data dependencies within the vector must be handled.

To work around these dependencies, Farrar (2006) introduces an alternative *striped* SIMD scheme where lanes are interleaved with each other. A*PA2 does not use this, but for example Shao and Ruan (2024) does.

Myers (1999) introduces a *bitpacking* algorithm specifically for edit distance (Fig. 1d). It bit-encodes the differences between *w* = 64 states in a column into two computer words and gives an efficient algorithm to operate on them. This provides a significant speedup over previous methods. The supplement of BitPAl (Loving *et al*., 2014; Benson *et al*., 2013) introduces an alternative scheme for edit distance based on a different bit-encoding, but as both methods end up using 20 instructions (see Appendix A.1) we did not pursue this further.

#### Tools

There are many semi-global aligners that implement *O*(*nm*) (semi)-global alignment using numerous of the aforementioned implementation techniques, such as SeqAn (Döring *et al*., 2008), Parasail (Daily, 2016), SWIPE (Rognes, 2011), Opal (Šošic, 2015), libssa (Frielingsdorf, 2015), SWPS3 (Szalkowski *et al*., 2008), SSW library (Zhao *et al*., 2013).

Dedicated global alignment implementations implementing band-doubling are much rarer. Edlib (Šošić and Šikić, 2017) implements *O*(*ns*) band doubling and Myers’ bitpacking (Fig. 1d). KSW2 implements band doubling for affine costs (Suzuki and Kasahara, 2018; Li, 2018). WFA and BiWFA (Marco-Sola *et al*., 2020, 2022) implement the *O*(*n* + *s*^2^) expected time *diagonal transition* algorithm (Ukkonen, 1985; Myers, 1986) (Fig. 1b). Block Aligner (Liu and Steinegger, 2023) is an approximate aligner that can handle position-specific scoring matrices whose main novelty is to divide the computation into larger blocks. Recently Shao and Ruan (2024) provided a new implementation of band doubling based on Farrar’s striped method that focusses on affine costs but also supports edit distance. Lastly, A*PA (Groot Koerkamp and Ivanov, 2024) directly implements A* on the alignment graph using the gap-chaining seed heuristic.

### 2 Preliminaries

#### Edit graph

We take as input two zero-indexed sequences *A* and *B* over an alphabet of size 4 of lengths *n* and *m*. The *edit graph* (Fig. 2) contains *states* ⟨*i, j*⟩ (0 ≤ *i* ≤ *n*, 0 ≤ *j* ≤ *m*) as vertices. It further contains directed insertion and deletion edges ⟨*i, j*⟩ → ⟨*i, j* + 1⟩ and ⟨*i, j*⟩ → ⟨*i* + 1, *j*⟩ of cost 1, and diagonal edges ⟨*i, j*⟩ → ⟨*i* + 1, *j* + 1⟩ of cost 0 when *A*_*i*_ = *B*_*i*_ and substitution cost 1 otherwise. The shortest path from *vs* := ⟨0, 0⟩ to *vt* := ⟨*n, m*⟩ in the edit graph corresponds to an optimal alignment of *A* and *B*. The *distance d*(*u, v*) from *u* to *v* is the length of the shortest (minimal cost) path from *u* to *v*, and we use *distance, length*, and *cost* interchangeably. We write *g*^∗^(*u*) := *d*(*vs, u*) for the distance from the start to *u, h*^∗^(*u*) := *d*(*u, vt*) for the distance from *u* to the end, and *f* ^∗^(*u*) := *g*^∗^(*u*) + *h*^∗^(*u*) for the minimal cost of a path through *u*.

**Fig. 2.**
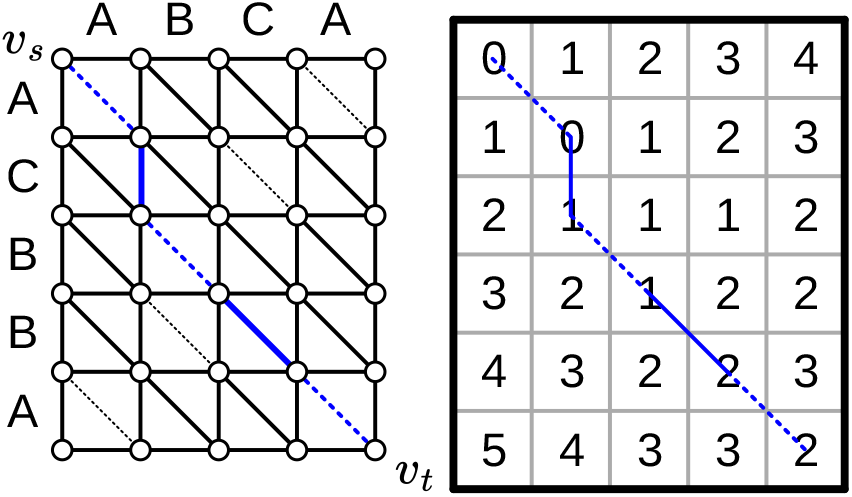
An example of an edit graph (left) corresponding to the alignment of strings `ABCA` and `ACBBA`, adapted from Sellers (1974). Solid edges indicate insertion/deletion/substitution edges of cost 1, while dashed edges indicate matches of cost 0. All edges are directed from the top-left to the bottom-right. The shortest path of cost 2 is shown in blue. The right shows the corresponding dynamic programming (DP) matrix containing the distance *g*^∗^(*u*) to each state.

**A\*** is a shortest path algorithm based on a *heuristic* function *h*(*u*) (Hart *et al*., 1968). A heuristic is called *admissible* when *h*(*u*) never overestimates the distance to the end, i.e., *h*(*u*) ≤ *h*^∗^(*u*), and admissible *h* guarantee that A* finds a shortest path. A* *expands* states in order of increasing *f* (*u*) := *g*(*u*) + *h*(*u*), where *g*(*u*) is the best distance to *u* found so far. We say that *u* is *fixed* when the distance to *u* has been found, i.e., *g*(*u*) = *g*^∗^(*u*).

#### Computational volumes

Spouge (1989) defines a *computational volume* as a subgraph of the alignment graph that contains all shortest paths. Given a bound *t* ≥ *s*, some examples of computational volumes are:

1. The entire (*n* + 1) × (*m* + 1) graph or DP table.
2. {*u* : *g*^∗^(*u*) ≤ *t*}, the states at distance ≤ *t*, introduced by Fickett (1984) and similar to Dijkstra’s algorithm (Figs. 1a and 1b).
3. {*u* : *c*gap(*vs, u*) + *c*gap(*u, vt*) ≤ *t*} the static set of states possibly on a path of cost ≤ *t* (Fig. 1c) (Ukkonen, 1985).
4. {*u* : *g*^∗^(*u*) + *c*gap(*u, vt*) ≤ *t*}, as used by Edlib (Figs. 1d and 1e) (Šošić and Šikić, 2017; Spouge, 1991).
5. {*u* : *g*^∗^(*u*) + *h*(*u*) ≤ *t*}, for any admissible heuristic *h*, which A*PA2 uses and is similar to A* (Figs. 1f and 1g).

**Band-doubling** is the following algorithm by Ukkonen (1985), that depends on the choice of computational volume being used.

1. Start with edit distance threshold *t* = 1.
2. Loop over columns *i* from 0 to *n*.
3. For each column, determine the range of rows [*j*_*start*_, *j*_*end*_] to be computed according to the computational volume that’s being used.
  a. If this range is empty or does not contain a state at distance ≤ *t*, double *t* and go back to step 1.
  b. Otherwise, compute the distance to the states in the range, and continue with the next column.

The algorithm stops when *t*_*k*_ ≥ *s > t*_*k*−1_. For the *c*_gap_(*v*_*s*_, *u*) + *c*gap(*u, vt*) ≤ *t* computational volume used by Ukkonen, each test requires *O*(*n* · *t*_*i*_) time, and hence the total time is

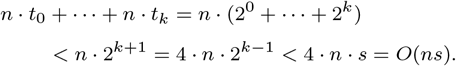

Note that with this computational volume we can not reuse values from previous iterations, resulting in roughly a factor 2 overhead.

**Myers’ bitpacking** exploits that the difference in distance *g*^∗^(*u*) to adjacent states is always in {−1, 0, +1} (Myers, 1999). The method bit-encodes *w* = 64 differences between adjacent states in a column in two indicator words, indicating positions where the difference is +1 and −1 respectively. Given also the similarly encoded difference along the top, a 1×*w* rectangle can be computed in only 20 bit operations (Appendix A.1). We call each consecutive non-overlapping chunk of 64 rows a *lane*, so that there are ⌈*m/*64⌉ lanes, where the last lane may be padded. Note that this method originally only uses 17 instructions, but some additional instructions are needed to support multiple lanes when *m > w*.

#### Profile

Instead of computing each substitution score *S*[*A*_*i*_][*B*_*j*_] = [*A*_*i*_ ≠ *B*_*j*_] for the 64 states in a word one by one, Myers’ algorithm first builds a *profile* (Rognes and Seeberg, 2000). For each character *c, Eq*[*c*] stores an ⌈*m/w*⌉ word bitvector indicating which characters of *B* equal *c*. This way, adjacent scores in a column are simply found as *Eq*[*A*_*i*_][*j* … *j*^*′*^].

**Edlib** implements band doubling using the *g*^∗^(*u*) + *c*gap(*u, vt*) ≤ *t* computational volume and bitpacking (Šošić and Šikić, 2017). For traceback, it uses Hirschberg’s *meet-in-the-middle* approach: once the distance is found, the alignment is started over from both sides towards the middle column, where a state on the shortest path is determined. This is recursively applied to the left and right halves until the sequences are short enough that *O*(*sn*) memory can be used.

## 3 Methods

Conceptually, A*PA2 builds on Edlib. First we describe how we make the implementation more efficient using SIMD and blocks. Then, we modify the algorithm itself by using a new traceback method and avoiding unnecessary recomputation of states. On top of that, we apply the A*PA heuristics for further speed gains on large and complex alignments, at the cost of larger precomputation time to build the heuristic.

### 3.1 Band-doubling

A*PA2 uses band-doubling with the *g*^∗^(*u*) + *h*(*u*) ≤ *t* computational volume. That is, in each *iteration* of *t* we compute the distance to all states with *g*^∗^(*u*) + *h*(*u*) ≤ *t*. In its simple form, we use *h*(*u*) = *c*gap(*u, vt*), like Edlib does. We start doubling at *h*(*vs*) = *h*(⟨0, 0⟩), so that *t*_*i*_ := *h*(⟨0, 0⟩) + *B* · 2^*i*^, where *B* is the block size introduced below.

### 3.2 Blocks

Instead of determining the range of rows to be computed for each column individually, we determine it once per *block* of *B* = 256 consecutive columns. This computes some extra states, but reduces the overhead by a lot. (From here on, *B* stands for the block size, and not for the sequence *B* to be aligned.) Within each block, we iterate over the *lanes* of *w* = 64 rows at a time, and for each lane compute all *B* columns before moving on to the next lane.

Section 3.8 explains in detail how the range of rows to be computed is determined.

### 3.3 Memory

Where Edlib does not initially store intermediate values and uses meet-in-the-middle to find the alignment, A*PA2 stores the distance to all states at the end of *each* block, encoded as the distance to the top-right state of the block and the bit-encoded vertical differences along the right-most column. This simplifies the traceback method (see Section 3.6), and has sufficiently small memory usage to be practical.

### 3.4 SIMD

While it is tempting to use a SIMD vector as a single *W* = 256-bit word, the four *w* = 64-bit words (SIMD lanes) are dependent on each other and require manual work to shift bits between the lanes. Instead, we let each 256-bit AVX2 SIMD vector represent four 64-bit words (lanes) that are anti-diagonally staggered as in Fig. 3a. This is similar to the original anti-diagonal tiling introduced by Wozniak (1997), but using units of *w*-bit words instead of single characters. This idea was already introduced in 2014 by the author of Edlib in a GitHub issue (github.com/Martinsos/edlib/issues/5), but to our knowledge has never been implemented in either Edlib or elsewhere.

**Fig. 3.**
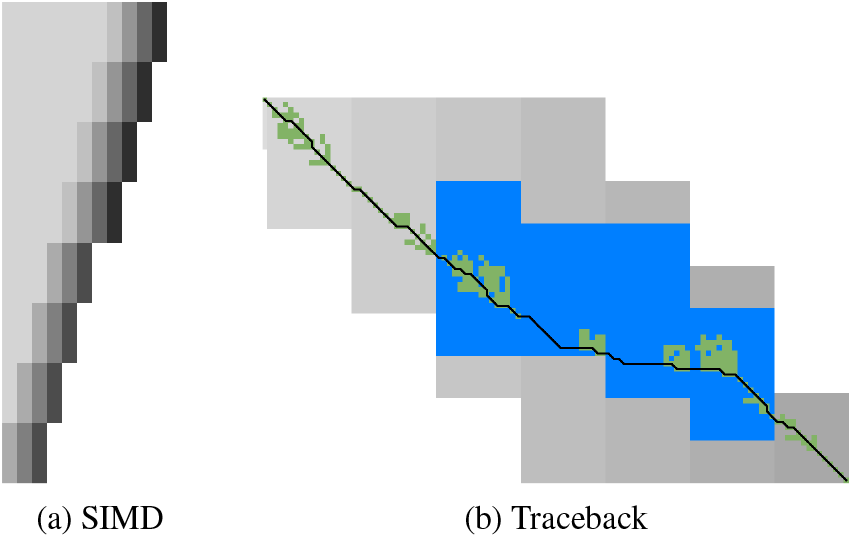
(a) SIMD processing of two times 4 lanes in parallel. This example uses lanes of 4 instead of 64 rows. First the top-left triangle is computed lane by lane, and then 8-lane diagonals are computed by using two 4-lane SIMD vectors in parallel. (b) States expanded by the diagonal transition traceback in each block are shown in green. When the distance in a block is too large, a part of the block is fully recomputed as fallback, as shown in blue. In regions with low divergence, diagonal transitoin is sufficient to trace the path, and only in noisy regions the algorithm falls back to recomputing full blocks.

We further improve instruction-level-parallelism (ILP) by processing 8 lanes at a time using two SIMD vectors in parallel, spanning a total of 512 rows (Fig. 3a).

When the number of remaining lanes to be computed is 𝓁, we process 8 lanes in parallel as long as 𝓁 ≥ 8. If there are remaining lanes, we end with another 8-lane (5 ≤ 𝓁 *<* 8) or 4-lane (1 ≤ 𝓁 ≤ 4) iteration that optionally includes some padding lanes at the bottom. In case the horizontal differences along the original bottom row are needed (as required by incremental doubling Section 3.9), we can not use padding and instead fall back to trying a 4-lane SIMD (𝓁 ≥ 4), a 2-lane SIMD (𝓁 ≥ 2), and lastly a scalar iteration (𝓁 ≥ 1).

### 3.5 SIMD-friendly sequence profile

A drawback of anti-diagonal tiling is that each lane corresponds to a column with character *a*_*i*_ that needs to be looked up in the profile *Eq*[*a*_*i*_][𝓁]. While SIMD can do multiple lookups in parallel using `gather` instructions, these instructions are not always efficient. Thus, we introduce the following alternative scheme. Let *b* = ⌈log_2_(*σ*)⌉ be the number of bits needed to encode each character, with *b* = 2 for DNA. For each lane, the new profile *Eq*^*′*^ stores *b* words as an ⌈*m/w*⌉ × *b* array *Eq*^*′*^[𝓁][*p*]. Each word 0 ≤ *p < b* stores the negation of the *p*th bit of each character. To check which characters in lane 𝓁 contain character *c* with bit representation 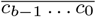, we precompute *b* words 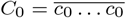 to 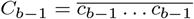 and then compute 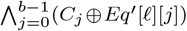, where ⊕ denotes the x or operation. As an example take *b* = 2 and a lane with *w* = 8 characters (0, 1, 2, 2, 3, 3, 3, 3). Then 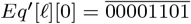 and 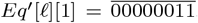, keeping in mind that bits are shown in reverse order in this notation. If the column now contains character 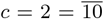 we initialize 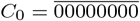 and 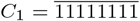 and compute

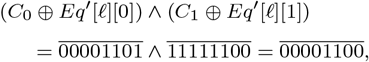

indicating that 0-based positions 2 and 3 contain character 2. This naturally extends to SIMD vectors, where each lane is initialized with its own constants.

### 3.6 Traceback

The traceback stage takes as input the computed vertical differences at the end of each block of columns. We iteratively work backwards through the blocks. When tracing the block covering columns *i* to *i* + *B*, we know the distances *g*(⟨*i, j*⟩) to the states in column *i* at the start of the block, and a state *u* = ⟨*i* + *B, j*⟩ at distance *g*^∗^(*u*) in column *i* + *B* at the end of the block that is on an optimal path. The goal is to find an optimal path from column *i* to *u*.

A naive approach is to simply recompute the entire block of columns while storing distances to all states. Here we consider two more efficient methods.

#### Optimistic block computation

Instead of computing the full range of rows for this column, a first insight is that only rows up to *j* are needed, since the optimal path to *u* = ⟨*i* + *B, j*⟩ can never go below row *j*.

Secondly, the path crosses *B* = 256 columns, and so we optimistically assume that it will be contained in rows *j* − 256 − 64 = *j* − 320 to *j*. Thus, we first recompute this range of rows (rounded out to multiples of *w* = 64) from left to right while storing intermediate values, as shown in blue in Fig. 3b. If the distance to *u* computed this way equals *g*^∗^(*u*), there is a shortest path contained within the computed rows and we trace it one state at a time. Otherwise, we repeatedly try again with double the number of lanes until success. The exponential search ensures low overhead and good average case performance.

#### Optimistic diagonal transition traceback (DTT)

A second improvement uses the *diagonal transition* algorithm backwards from *u*. We simply run the unmodified algorithm on the reverse graph covering columns *i* to *i* + *B* and rows 0 to *j*. Whenever a state *v* in column *i* is reached, say at distance *d* from *u*, we check whether *g*(*v*) + *d* = *g*^∗^(*u*), and continue until a *v* is found for which this holds. We then know that *v* lies on a shortest path and can find the path from *v* to *u* via a usual traceback on the diagonal transition algorithm, as shown in green in Fig. 3b.

As an optimization, when no suitable *v* is found after trying all states at distance ≤ 40, we abort the DTT and fall back to the block doubling described above. Another optimization is the WF-adaptive heuristic introduced by WFA: all states that lag more than 10 behind the furthest reaching diagonal are dropped. Lastly, we abort early when after reaching distance 20 = 40*/*2, less than half the columns of the block were traversed.

### 3.7 A*

Edlib already uses a simple *gap-cost* heuristic that gives a lower bound on the number of insertions and deletions on a path from each state to the end. We replace this by the much stronger gap-chaining seed heuristic (GCSH) introduced in A*PA, with two modifications.

#### Contours update

In A*PA, matches are *pruned* as soon as a shortest path to their start has been found. This helps to penalize states *before* (left of) the match. Each iteration of A*PA2 works left-to-right only, and thus pruning of matches does not affect the current iteration. Thus, we collect all matches to be pruned at the end of each iteration, and update the contours in one right-to-left sweep.

To keep the computational volume valid after pruning, we ensure that the range of computed rows never shrinks.

#### Pre-pruning

Here we introduce an independent optimization that also applies to the original A*PA method. Each of the heuristics *h* introduced in A*PA depends on the set of *matches ℳ*. First the sequence *A* is split into *seeds s*_*i*_ of length *k*, and then *ℳ* consists of the exact matches in sequence *B* of all seeds. When *ℳ* contains *all* matches, *h* is an admissible heuristic that never overestimates the true distance. And it was shown that even after pruning some matches, *h* is still a lower bound on the length of any path not going through already expanded states.

Now consider an exact match *m* for seed *s*_*i*_ from *u* to *v*. The existence of the match is a ‘promise’ that seed *s*_*i*_ can be crossed for free (cost less than *r* = 1). When *m* is a match outside the optimal alignment, it is likely that *m* can not be extended into a longer alignment. When indeed *m* can not be extended into an alignment of *s*_*i*_ and *s*_*i*+1_ of cost less than 2, the existence of *m* was a ‘false promise’, since crossing the two seeds takes cost at least 2. Thus, we can ignore *m* and remove *m* from the heuristic, making the heuristic more accurate.

More generally, we try to extend each match *m* into an alignment covering seeds *s*_*i*_ up to (but excluding) *s*_*i*+*q*_ for all *q* ≤ *p* = 14. If any of these extensions has cost at least *q*, i.e., *m* falsely promised that *s*_*i*_ to *s*_*i*+*q*_ can be crossed for cost *< q*, we *pre-prune* (remove) *m*.

We try to extend each match by running the diagonal transition algorithm from the end of each match, and dropping any furthest reaching points that are at distance ≥ *q* while at most *q* seeds have been covered.

As shown in Fig. 4b, the effect is that the number of off-path matches is significantly reduced. This makes contours simpler and hence faster to initialize, update, and query, and it increases the value of the heuristic

**Fig. 4.**
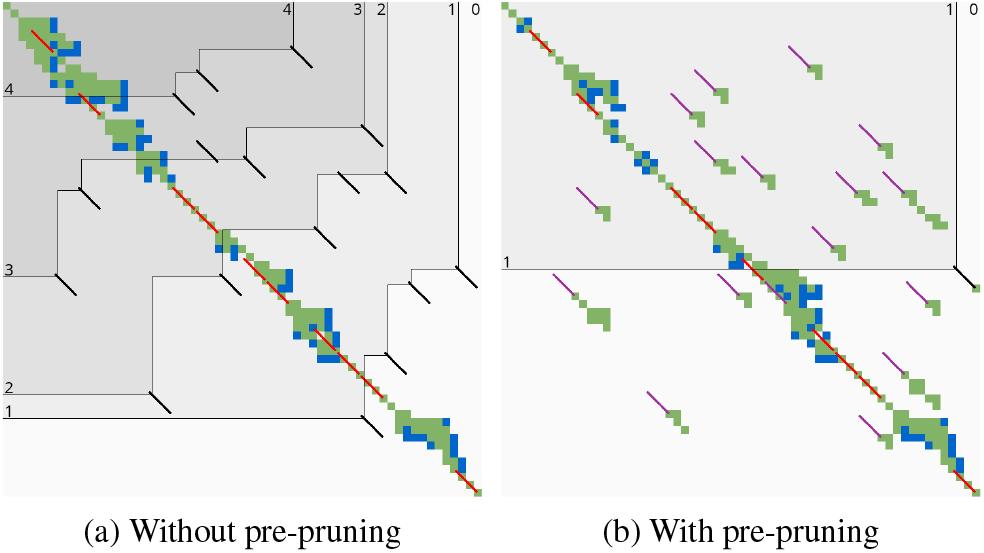
Effect of pre-pruning on chaining seed heuristic (CSH) contours. The left shows contours and layers of the heuristic at the end of an A*PA alignment, after matches (black diagonals) on the path have been pruned (red diagonal). The right shows pre-pruned matches in purple and the states visited during pre-pruning in green. After pre-pruning, almost no off-path matches remain. This decreases the number of contours, making the heuristic stronger, and simplifies contours, making the heuristic faster to evaluate.

### 3.8 Determining the rows to compute

For each block, spanning columns *i* to *i* + *B*, only a subset of rows is computed in each iteration. Namely, we only compute those rows that can possibly contain states on a path/alignment of cost at most *t*. Intuitively, we try to ‘trap’ the alignment inside a wall of states that can not lie on a path of length at most *t*, i.e., have *f* ^∗^(*u*) ≥ *t*, as can be seen in Fig. 5a. We determine this range of rows in two steps:

**Fig. 5.**
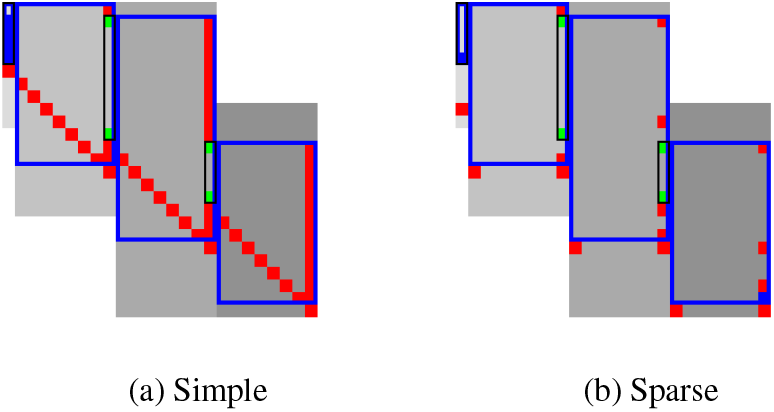
Detail of computed ranges. Coloured states are invocations of *f* . Red: *f* (*u*) *> t* or *f*_*l*_(*u*) *> t*, green: *f* (*u*) ≤ *t* and *u* is fixed, and blue: *f*_*l*_(*u*) ≤ *t*. Vertical black rectangles indicated fixed states, and blue rectangles indicate the range of rows 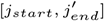 that must be computed for each block. The third block has no fixed states in its right column, indicating that *t* must be increased.

1. First, we determine the *fixed range* at the end of the preceding block. I.e., we find the topmost and bottommost states ⟨*i, jstart*⟩ and ⟨*i, j*_*end*_⟩ with *f* (*u*) = *g*(*u*) + *h*(*u*) ≤ *t*, as shown in green in Fig. 5. All in-between states *u* = ⟨*i, j*⟩ with *j*_*start*_ ≤ *j* ≤ *j*_*end*_ are then *fixed*, meaning that the correct distance has been found and *g*(*u*) = *g*^∗^(*u*).

2. Then, we use the heuristic to find the bottommost state 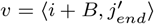 at the end of the to-be-computed block that can possibly lie on a path of length ≤ *t*. We then compute rows *jstart* to 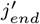 in columns *i* to *i* + *B*, rounding 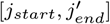 *out* to the previous/next multiple of the word size *w* = 64.

**Step 1: Fixed range**. Suppose that states in rows [*r*_*start*_, *r*_*end*_] were computed. One way to find *j*_*start*_ and *j*_*end*_ is by simply iterating inward from the start/end of the range and skipping all states with *f* (*u*) = *g*(*u*)+ *h*(*u*) *> t*, as indicated by the red columns in Fig. 5a.

**Step 2: End of computed range**. We will now determine the bottommost row *j* that can contain a state on a path of length ≤ *t* at the end of the block. Let *u* = ⟨*i, j*_*end*_⟩ be the bottommost fixed state in column *i* with *f* (*u*) ≤ *t*. Let *v* = ⟨*i*^*′*^, *j*^*′*^⟩ be a state in the current block (*i* ≤ *i*^*′*^ ≤ *i* + *B*) that is below the diagonal of *u*. Suppose *v* lies on a path of length ≤ *t*. This path most cross column *i* in or above *u*, since states *u*^*′*^ below *u* have *f* ^∗^(*u*^*′*^) *> t*. The distance to *v* is now at least min_*j*≤*jend*_*g*^∗^(⟨*i, j*⟩) + *c*gap(⟨*i, j*⟩, *v*) ≥ *g*^∗^(*u*) + *c*gap(*u, v*), and thus we define

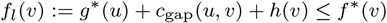

as a lower bound on the length of the shortest path through *v*, assuming *v* is below the diagonal of *u* and *f* ^∗^(*v*) ≤ *t*. When *f*_*l*_(*v*) *> t*, this implies *f* ^∗^(*v*) *> t* and also *f* ^∗^(*v*^*′*^) *> t* for all *v*^*′*^ below *v*. The end of the range is now computed by finding the bottommost state *v* in each column for which *f*_*l*_ is at most *t*, using the following algorithm:

1. Start with *v* = ⟨*i*^*′*^, *j*^*′*^⟩ = *u* = ⟨*i, j*_*end*_⟩.

2. While the below-neighbour *v*^*′*^ = ⟨*i*^*′*^, *j*^*′*^ + 1⟩ of *v* has *f*_*l*_(*v*) ≤ *t*, increment *j*^*′*^.

3. Go to the next column by incrementing *i*^*′*^ and *j*^*′*^ by 1 and repeat step 2, until *i*^*′*^ = *i* + *B*.

The row 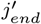 of the last *v* we find in this way is the bottommost state in column *i* + *B* that can possibly have *f* (*v*) ≤ *t*, and hence this is end of the range we compute.

In Fig. 5a, we see that *f* (*v*) is evaluated at a diagonal of states just below the bottommost fixed (green) state *u* at the end of the preceding black, and that the to-be-computed range (indicated in blue) includes exactly all states above this diagonal.

**Sparse heuristic invocation**. A drawback of the previous method is that it requires a large number of calls to *f* and hence the heuristic *h*: roughly one per column and one per row. In Appendix A.2 we present a *sparse* version that uses fewer calls to *f*, as shown in Fig. 5b.

### 3.9 Incremental doubling

When the original band doubling algorithm doubles the threshold from *t* to 2*t*, it simply recomputes the distance to all states. On the other hand, BFS, Dijkstra, and A* with a consistent heuristic visit states in increasing order of distance (*g*(*u*) for BFS and Dijkstra, *f* (*u*) = *g*(*u*) + *h*(*u*) for A*), and the distance to a state is known to be correct (*fixed*) as soon as it is expanded for the first time. This way a state is never expanded twice.

Indeed, our band-doubling algorithm can also avoid recomputations. After completing the iteration for *t*, it is guaranteed that the distance is fixed to all states that indeed satisfy *f* (*u*) ≤ *t*. In fact a stronger result holds: in any column the distance is fixed for all states *between* the topmost and bottommost state in that column with *f* (*u*) ≤ *t*.

To be able to skip rows, we must store horizontal differences along a row so we can continue from there in the next iteration. We choose this row *j*_*f*_ (for *fixed*) as the last row at a lane boundary before the end of the fixed states in the last column of the preceding block, as indicated in Fig. 6 by a horizontal black rectangle. In the first iteration, reusing values is not possible, so we split the computation of the block into two parts (Fig. 6a): one above *j*_*f*_, to extract and store the horizontal differences at *j*_*f*_, and the remainder below *j*_*f*_ .

**Fig. 6.**
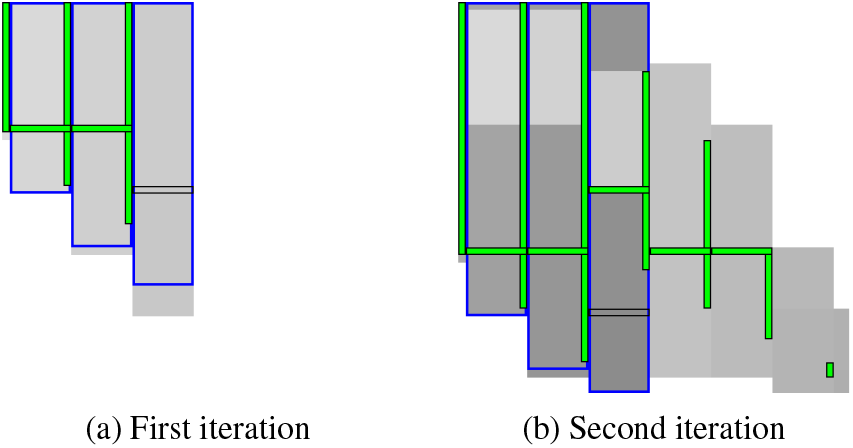
Incremental doubling detail. Blue rectangles show the ranges required to be computed, and grey the computed blocks. Vertical green rectangles show the fixed range at the end of each block, and green horizontal rectangles a fixed row *j*_*f*_ of states inside some blocks. In both figures the third block was just computed, in the first (left) and second (right) iteration of trying a threshold. The black horizontal rectangle indicates the new candidate 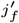 for the fixed horizontal region.

In the second and further iterations, the values at *j*_*f*_ are reused and the block is split into three parts. The first part computes all lanes covering states before the start of the fixed range at the end of the block (the green column at the end of the third column in Fig. 6b). We skip the lanes up to the previous *j*_*f*_, since the values at both the bottom and right of this region are already fixed. Then, we compute the lanes between the old *j*_*f*_ and its new value 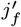 . Lastly we compute the lanes from 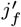 to the end.

## 4 Results

Our implementation A*PA2 is written in Rust and available at github.com/RagnarGrootKoerkamp/astar-pairwise-aligner. We compare it against other aligners on real datasets, report the impact of the individual techniques we introduced, and measure time and memory usage.

### 4.1 Setup

#### Datasets

We benchmark on five datasets containing real sequences of varying length and divergence, as listed in detail in Table 1 in Appendix A.3. They can be downloaded from github.com/pairwise-alignment/pa-bench/releases/tag/datasets.

**Table 1.** Dataset statistics. All but the first dataset are ONT reads. The dataset with genetic variation (gv) also includes long gaps, while the SARS-CoV-2 dataset stands out for having only 1.5% divergence on average. **Cnt**: number of sequence pairs. **Max gap**: longest gap in the reconstructed alignment.

| Dataset | Source | Cnt | Length [kbp] |  |  | Divergence [%] |  |  | Max gap [kbp] |  |
| --- | --- | --- | --- | --- | --- | --- | --- | --- | --- | --- |
|  |  |  | min | mean | max | min | mean | max | mean | max |
| SARS-CoV-2 | A*PA2 | 10 000 | 27 | 30 | 30 | 0.0 | **1.5** | 12.8 | 0.1 | 1.0 |
| ONT 1 kbp | WFA | 12 500 | 0.04 | 0.8 | 1.1 | 0.0 | 10.4 | 22.5 | 0.01 | 0.1 |
| ONT 10 kbp | BiWFA | 10 000 | 0.2 | 11 | 50 | 3.0 | 11.6 | 19.2 | 0.07 | 3.4 |
| ONT >500 kbp | A*PA | 50 | 500 | 594 | 849 | 2.7 | 6.1 | 16.7 | 0.1 | 1.3 |
| ONT >500 kbp + gv | BiWFA | 48 | 502 | 632 | 1053 | 4.3 | 7.2 | 18.2 | **1.9** | 42 |

Four datasets containing Oxford Nanopore Technologies (ONT) reads are reused from the WFA, BiWFA, and A*PA evaluations (Marco-Sola *et al*., 2020, 2022; Groot Koerkamp and Ivanov, 2024). Of these, two ‘*>*500 kbp’ and ‘*>*500 kbp with genetic variation’ datasets have divergence 6 − 7%, while two ‘1 kbp’ and ‘10 kbp’ datasets are filtered for sequences of length *<*1 kbp and *<*50 kbp and have average divergence 11% and average sequence length 800 bp and 11 kbp.

A SARS-CoV-2 dataset was newly generated by downloading 500 MB of viral sequences from the COVID-19 Data Portal, covid19dataportal.org (Harrison *et al*., 2021), filtering out non-ACTG characters, and selecting 10000 random pairs. This dataset has average divergence 1.5% and average length 30 kbp.

For each set, we sorted all sequence pairs by edit distance and split them into 50 files each containing multiple pairs, so that the first file contains the 2% of pairs with the lowest divergence. Reported runtimes are averaged over the sequences in each file.

#### Algorithms and aligners

We benchmark A*PA2 against state-of-the-art exact aligners Edlib, BiWFA, and A*PA. We further compare against the approximate aligners WFA-Adaptive (Marco-Sola *et al*., 2020) and Block Aligner. For WFA-Adaptive we use default parameters (10, 50, 10), dropping states that lag behind by more than 50. For Block Aligner we use block sizes from 0.1% to 1% of the input size. Block Aligner only supports affine costs so we use gap-open cost 1 instead of 0.

We compare two versions of A*PA2. A*PA2*-simple* uses all engineering optimizations (bitpacking, SIMD, blocks, new traceback) and uses the simple gap-heuristic. A*PA2*-full* additionally uses more complicated techniques: incremental-doubling, and the gap-chaining seed heuristic introduced by A*PA with pre-pruning.

#### Parameters

For A*PA2, we fix block size *B* = 256. For A*PA2-full, we use the gap-chaining seed heuristic (GCSH) of A*PA with exact matches (*r* = 1) and seed length *k* = 12. We pre-prune matches by looking ahead up to *p* = 14 seeds. A detailed parameter comparison can be found in Appendix A.3. For A*PA, we use inexact matches (*r* = 2) with seed length *k* = 15 by default, and only change this for the low-divergence SARS-CoV-2 dataset and 4% divergence synthetic data, where we use exact matches (*r* = 1) instead.

#### Execution

We ran all benchmarks using PaBench (github.com/pairwise-alignment/pa-bench) on Arch Linux on an `Intel Core i7-10750H` with 64GB of memory and 6 cores, with hyper-threading disabled, frequency boost disabled, and CPU power saving features disabled. The CPU frequency is fixed to 3.6GHz and we run 1 single-threaded job at a time with niceness −20. Reported running times are the average wall-clock time per alignment and do not include the time to read data from disk. For A*PA2-full, reported times do include the time to find matches and initialize the heuristic.

### 4.2 Comparison with other aligners

#### Speedup on real data

Fig. 7 compares the running time of aligners on real datasets. Table 2 in Appendix A.3 contains a corresponding table of average runtimes. On the *>*500 kbp ONT reads, A*PA2-full is 19× faster than Edlib, BiWFA, and A*PA in average running time, and using the gap-chaining seed heuristic in A*PA2-full provides 3× speedup over A*PA2-simple.

**Table 2.** **Average runtime per sequence** of each aligner on each dataset. Cells marked with *>* are a lower bound due to timeouts. Speedup is reported as the fastest A*PA2 variant compared to the fastest of Edlib, BiWFA, and A*PA. WFA-Adaptive and Block Aligner are approximate aligners.

| Aligner | SARS-CoV-2 | ONT |  |  |  |
| --- | --- | --- | --- | --- | --- |
|  | [ms] | 1 kbp [ms] | 10 kbp [ms] | >500 kbp [s] | >500 kbp + gv [s] |
| EDLIB | 11.14 | 0.110 | 8.0 | 3.74 | 5.20 |
| BiWFA | 1.13 | 0.042 | 9.3 | 4.47 | 6.96 |
| A*PA | 6.25 | 0.514 | >190.1 | >14.01 | >12.92 |
| WFA-Adaptive | 0.85 | 0.038 | 3.0 | 1.04 | 0.81 |
| BLOCK ALIGNER | 2.35 | 0.038 | 0.9 | 0.63 | 0.68 |
| A*PA2-simple | 0.89 | 0.052 | 1.4 | 0.58 | 0.78 |
| A*PA2-full | 2.00 | 0.083 | 1.7 | 0.20 | 0.27 |
| Speedup [×] | 1.3 | 0.81 | 5.6 | 18.8 | 19.0 |

**Fig. 7.**
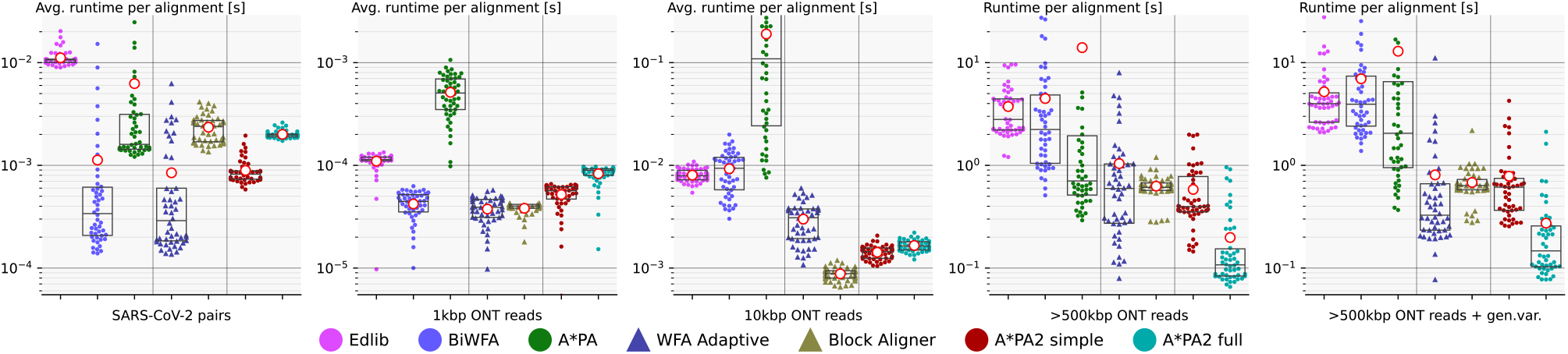
Runtime comparison (log). Each dot shows the running time of a single alignment (right two plots) or the average runtime over 2% of the input pairs (left four plots). Box plots show the three quartiles, and the red circled dot shows the average running time over all alignments. For A*PA, exact matches (*r* = 1) are used for the SARS-CoV-2 dataset, some alignments ≥ 10 kbp time out, and the shown average is a lower bound on the true average. Approximate aligners WFA Adaptive and Block Aligner are indicated with a triangle. On the *>*500 kbp reads, A*PA2-full is 19*×* faster than other exact methods.

On shorter sequences, the overhead of initializing the heuristic in A*PA2-full is large, and A*PA2-simple is faster. For the 10 kbp dataset, A*PA2-simple is 5.6× faster than other exact methods. For the shortest (1 kbp ONT reads) and most similar sequences (SARS-CoV-2 with 1% divergence), BiWFA is usually faster than Edlib and A*PA2-simple. In these cases, the overhead of using 256 wide blocks is relatively large compared to the edit distance *s* ≤ 500 in combination with BiWFA’s *O*(*s*^2^ + *n*) expected running time.

#### Comparison with approximate aligners

For the smallest datasets, BiWFA is about as fast as the approximate methods WFA Adaptive and Block Aligner, while for the largest datasets A*PA2-full is significantly faster. Only on the dataset of 10 kbp ONT reads is Block Aligner 1.6× faster than A*PA2, but it only finds the correct edit distance for 53% of the alignments. All accuracy numbers can be found in Table 3 in Appendix A.3.

**Table 3.**
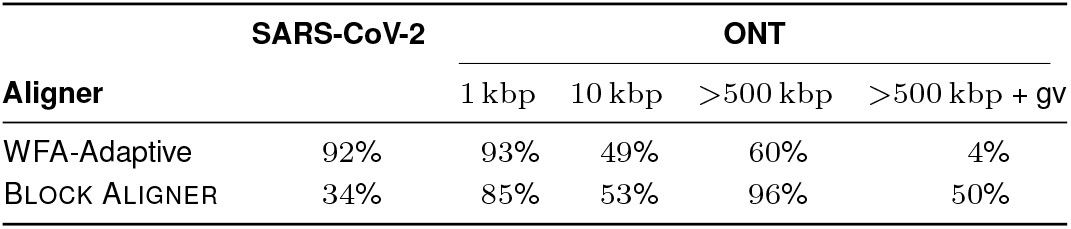
**Percentage of correctly aligned reads** by approximate aligners. The accuracy of WFA-Adaptive drops a lot for the *>*500 kbp dataset with genetic variation, since these alignments contain gaps of thousands of basepairs, much larger than the 50 bp cutoff after which trailing diagonals are dropped.

| Aligner | SARS-CoV-2 | ONT |  |  |  |
| --- | --- | --- | --- | --- | --- |
|  |  | 1 kbp | 10 kbp | >500 kbp | >500 kbp + gv |
| WFA-Adaptive | 92% | 93% | 49% | 60% | 4% |
| BLOCK ALIGNER | 34% | 85% | 53% | 96% | 50% |

#### Scaling with length

Figs. 8a and 8b compare the runtime of aligners on synthetic random sequences of increasing length and constant uniform divergence. BiWFA’s runtime is quadratic and is fast for sequences up to 3000 bp. As expected, A*PA2-simple has very similar scaling to Edlib but is faster by a constant factor. A*PA2-full includes the gap-chaining seed heuristic used by A*PA, resulting in comparable speed and near-linear scaling for both of them when *d* = 4%. For more divergent sequences, A*PA2-full is faster than A*PA since initializing the A*PA heuristic with inexact matches is relatively slow. The reason A*PA2-full is slower than A*PA for sequences of length 10 Mbp is that A*PA2-full uses seed length *k* = 12 instead of *k* = 15, causing the number of matches to explode when *n* approaches 4^12^ ≈ 16 · 10^6^.

**Fig. 8.**
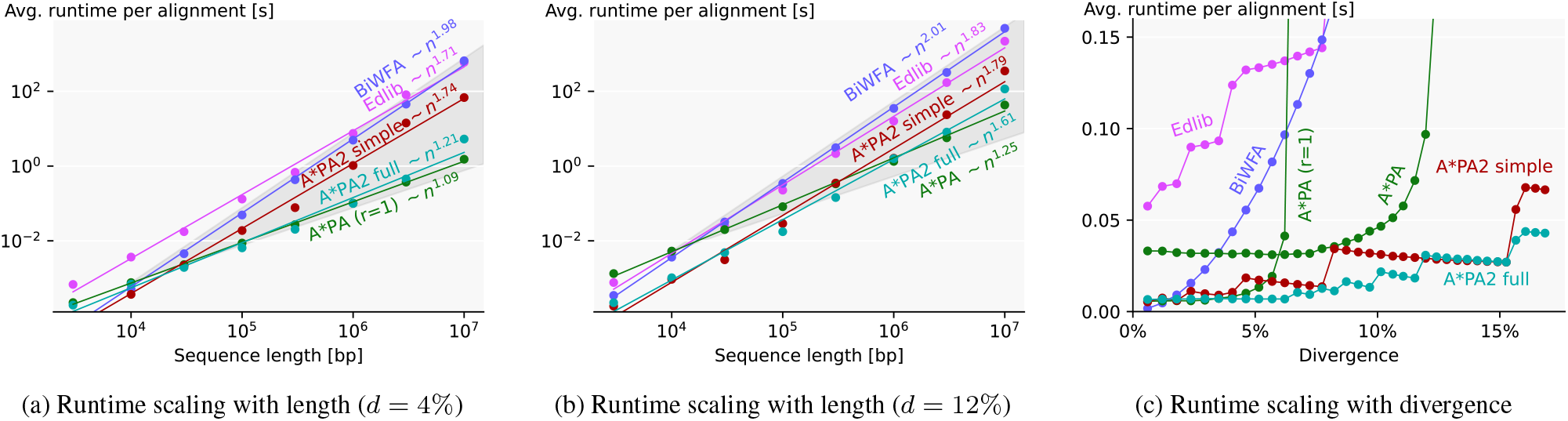
Runtime comparison on synthetic data. (a)(b) Log-log plot of average running time of aligners on synthetic sequences of increasing length with 4% divergence and 12% divergence. A*PA uses exact matches (*r* = 1) for *d* = 4% and inexact matches (*r* = 2) for *d* = 12%. For sequences of length *n*, averages are over 10^7^ */n* pairs. Lines are fitted in the log-log domain. The region between linear and quadratic growth is shaded in grey. (c) Average running time of aligners over 10 sequences of length 100 kbp with varying uniform divergence.

#### Scaling with divergence

Fig. 8c compares the runtime of aligners on synthetic sequences of increasing divergence. BiWFA’s runtime grows quadratically, while Edlib grows linearly and jumps up each time another doubling of the threshold is required. A*PA is fast until the maximum potential is reached at 6% resp. 12% and then becomes very slow. A*PA2 behaves similar to Edlib and jumps up each time another doubling of the threshold is needed, but is much faster. It outperforms BiWFA for divergence ≥ 2% and A*PA for divergence ≥ 4%. The runtime of A*PA2-full is near-constant up to divergence 7% due to the gap-chaining seed heuristic which can correct for up to 1*/k* = 1*/*12 = 8.3% of divergence, while A*PA2-simple starts to slow down at lower divergence. For a fixed number of doublings of the threshold, A*PA2 is faster for higher divergence because too low thresholds are rejected more quickly.

**Memory usage** of A*PA2 on *>*500 kbp sequences is at most 200 MB and only 30 MB in median. On all other datasets, memory usage is always less than 10 MB (Table 4 in Appendix A.3).

**Table 4.** **Memory usage** of aligners, measured as the increase in `max_rss` before and after aligning a pair of sequences.

| Aligner | SARS-CoV-2 |  | ONT |  |  |  |  |  |  |  |
| --- | --- | --- | --- | --- | --- | --- | --- | --- | --- | --- |
|  |  |  | 1 kbp |  | 10 kbp |  | >500 kbp |  | >500 kbp + gv |  |
| Memory [MB] | median | max | median | max | median | max | median | max | median | max |
| EDLIB | 0 | 0 | 0 | 0 | 0 | 0 | 0 | 0 | 0 | 0 |
| BiWFA | 0 | 0 | 0 | 0 | 0 | 0 | 4 | 11 | 0 | 2 |
| A*PA | 0 | 236 | 0 | 0 | 228 | 873 | 84 | 3453 | 158 | 6868 |
| WFA-Adaptive | 0 | 11 | 0 | 0 | 0 | 0 | 0 | 0 | 0 | 0 |
| BLOCK ALIGNER | 0 | 16 | 0 | 0 | 0 | 3 | 583 | 1189 | 610 | 2171 |
| A*PA2 simple | 2 | 5 | 0 | 0 | 4 | 6 | 0 | 55 | 2 | 164 |
| A*PA2 full | 0 | 0 | 0 | 0 | 0 | 0 | 30 | 82 | 6 | 141 |

### 4.3 Effects of methods

Fig. 9 shows the effect of one-by-one adding improvements to A*PA2 on *>*500 kbp long sequences, starting with Ukkonen’s band-doubling method using Myers’ bitpacking. We first change to the *g*^∗^(*u*) + *c*_gap_(*u, vt*) computational volume, making it comparable to Edlib. Then we process blocks of 256 columns at a time and only store differences at block boundaries, giving 2.5× speedup. Adding SIMD gives another 2.7× speedup, and instruction level parallelism (ILP) provides a further small improvement. The diagonal transition traceback (DTT) and sparse heuristic computation do not improve performance of A*PA2-simple much on long sequences, but their removal can be seen to slow it down for shorter sequences in an ablation (Fig. 12 in Appendix A.3). Incremental doubling (ID), the gap-chaining seed heuristic (GCSH), pre-pruning (PP), and the pruning of A*PA give another 3× speedup on average and 4× speedup in the first quantile.

**Fig. 9.**
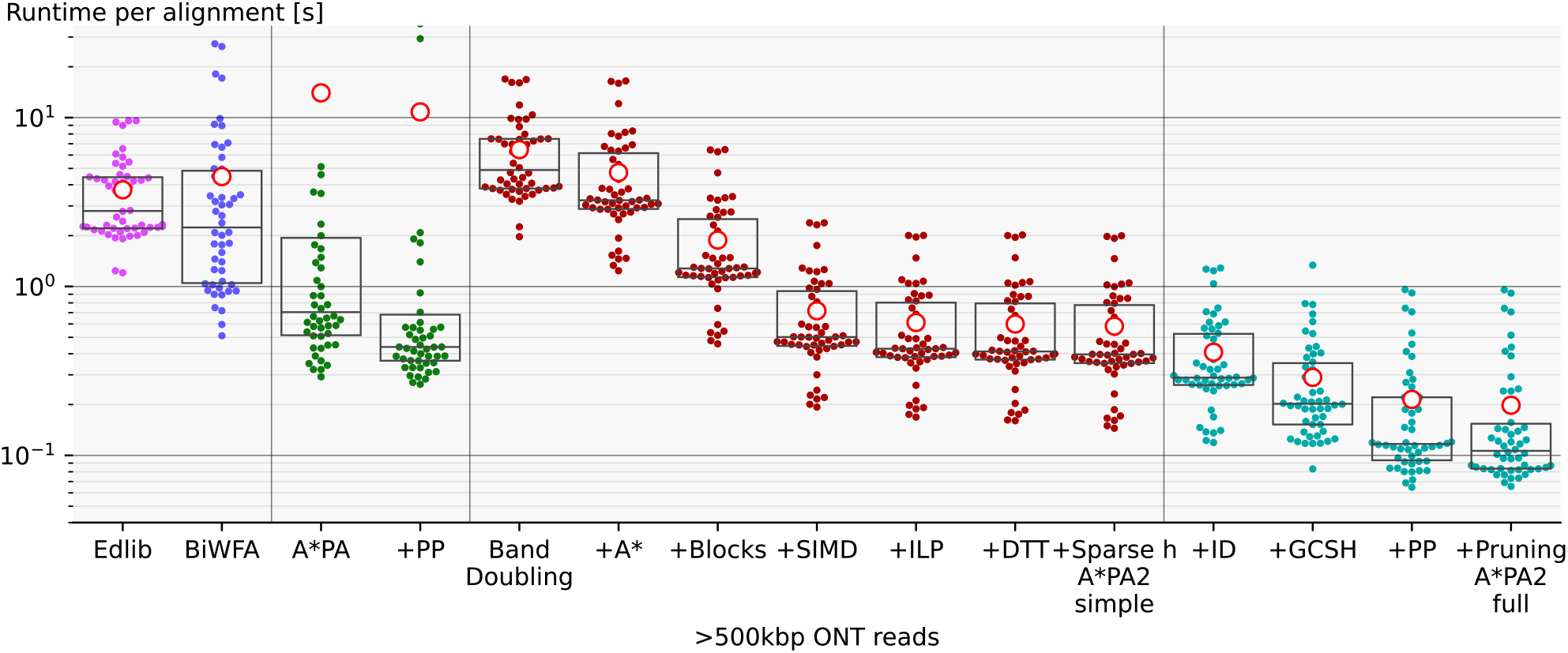
Effect of adding features. Box plots showing the performance improvements of A*PA2 when incrementally adding new methods one-by-one. A*PA2-simple corresponds to the rightmost red columns, and A*PA2-full corresponds to the rightmost blue column.

Fig. 14 in Appendix A.3 shows that A*PA2 spends most of its time computing blocks. For short 1 kbp long sequences, half the time is spent on traceback, and for the *>*500 kbp sequences, A*PA2-full spends around a quarter of time on initializing the heuristic.

## 5 Discussion

We have shown that by incorporating many existing techniques and by writing highly performant code, A*PA2 achieves 19× speedup over other methods when aligning *>*500 kbp ONT reads with 6% divergence, 5.6× speedup for sequences of average length 11 kbp, and only a slight slowdown over BiWFA for very short (*<*1 kbp) and very similar (*<*2% divergence) sequences. A*PA2’s speed is also comparable to approximate aligners, and is faster for long sequences, thereby nearly closing the gap between approximate and exact methods. A*PA2 achieves this by building on Edlib, using band doubling, bitpacking, blocks, SIMD, the gap-chaining seed heuristic, and pre-pruning. The effect of this is that A*PA2-simple has similar scaling behaviour as Edlib in both length and divergence, but with a significantly better constant. A*PA2-full additionally includes the A*PA heuristics and achieves the best of both worlds: the near-linear scaling with length of A*PA when divergence is small, and the efficiency of Edlib.

### Limitations

1. The main limitation of A*PA2-full is that the heuristic requires finding all matches between the two input sequences, which can take long compared to the alignment itself.
2. For sequences with divergence *<*2%, BiWFA exploits the sparse structure of the diagonal transition algorithm. In comparison, computing full blocks of size around 256 × 256 in A*PA2 has considerable overhead.
3. Only sequences over alphabet size 4 are currently supported, so DNA sequences containing e.g. N characters must be cleaned first.

### Future work

1. When divergence is low, performance could be improved by applying A* to the diagonal transition algorithm directly, instead of using DP. As a middle ground, it may be possible to compute individual blocks using diagonal transition when the divergence is low.
2. Currently A*PA2 is completely unaware of the type of sequences it aligns. Using an upper bound on the edit distance, either known or found using a non-exact method, could avoid trying overly large thresholds and smoothen the curve in Fig. 8c.
3. It should be possible to extend A*PA2 to open-ended and semi-global alignment, just like Edlib and WFA support these modes. Sequence-to-graph alignment may also be possible, but will likely be significantly more involved.
4. Extending A*PA2 to affine cost models should also be possible. This will require adjusting the gap-chaining seed heuristic, and changing the computation of the blocks from a bitpacking approach to one of the SIMD-based methods for affine costs.
5. Lastly, TALCO (Tiling ALignment using COnvergence of traceback pointers, https://turakhia.ucsd.edu/research/) provides an interesting idea: it may be possible start traceback while still computing blocks, thereby saving memory.

## Acknowledgements

I am grateful to Daniel Liu for discussions, feedback, and suggesting additional related papers, to André Kahles, Harun Mustafa, and Gunnar Rätsch for feedback on the manuscript, to Andrea Guarracino and Santiago Marco-Sola for sharing the WFA and BiWFA benchmark datasets, and to Gary Benson for help with debugging the BITPAL bitpacking code.

## Data availability

The data underlying this article are available on GitHub at github.com/pairwise-alignment/pa-bench/releases/tag/datasets

## A Appendix

### A.1 Bitpacking

Fig. 10 shows a SIMD version of Myers’ bitpacking and Fig. 11 shows the method for edit distance described in the supplement of Loving *et al*. (2014). Both methods require 20 instructions.

**Fig. 10.**
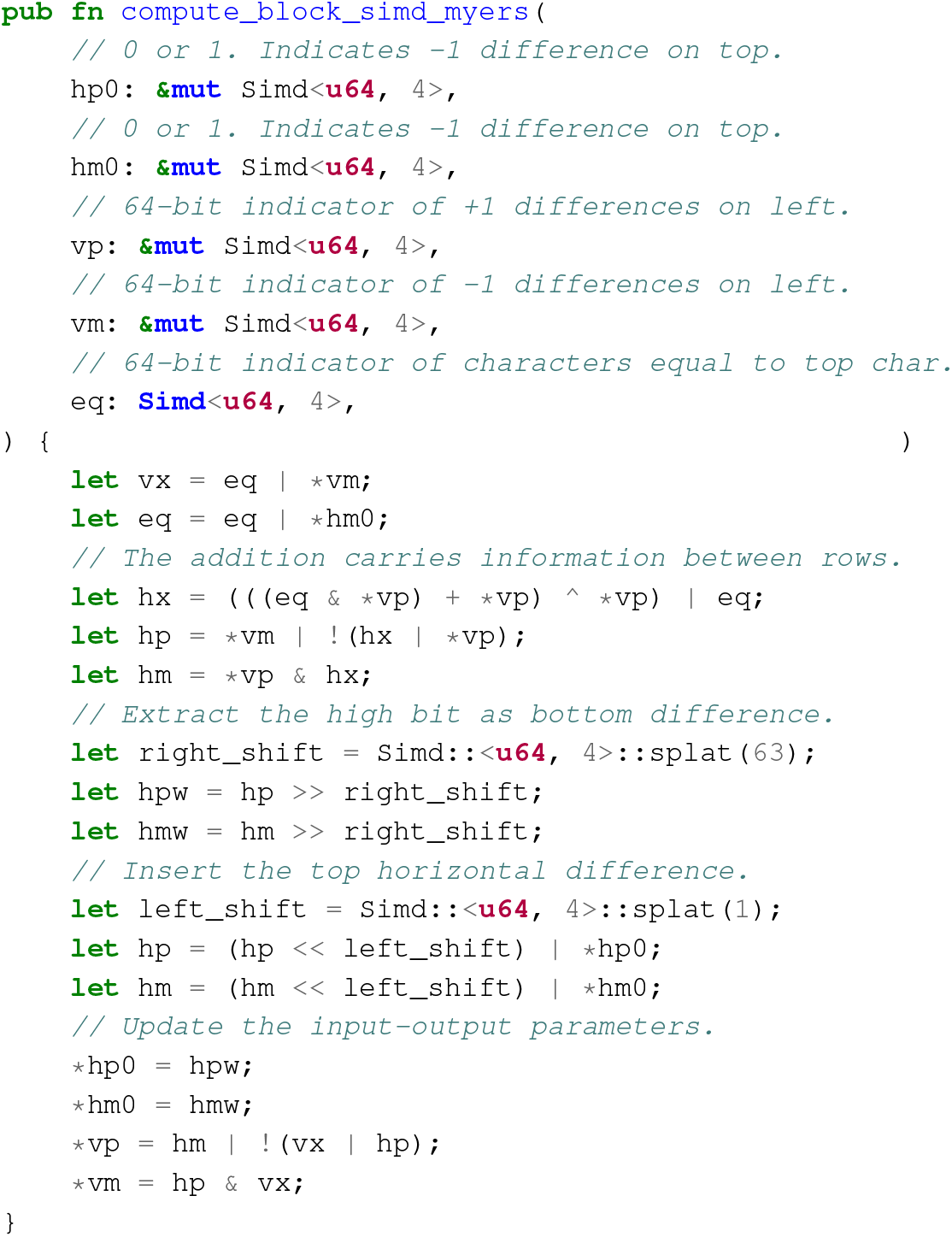
Myers’ bitpacking. Rust code for SIMD version of Myers’ bitpacking algorithm that takes 20 instructions.

**Fig. 11.**
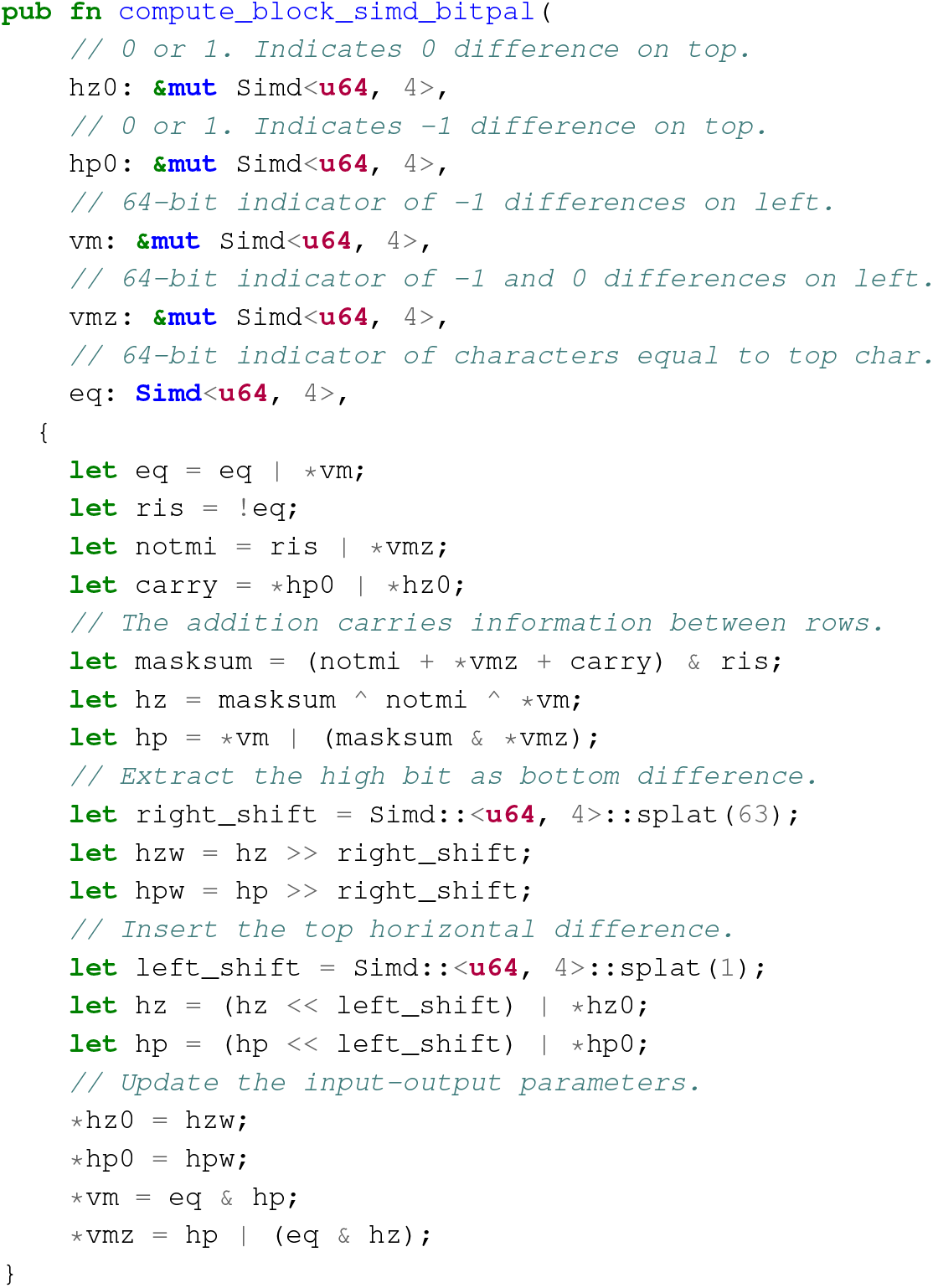
BITPAL’s bitpacking. Rust code for SIMD version of BITPAL’s bitpacking algorithm that takes 20 instructions.

**Fig. 12.**
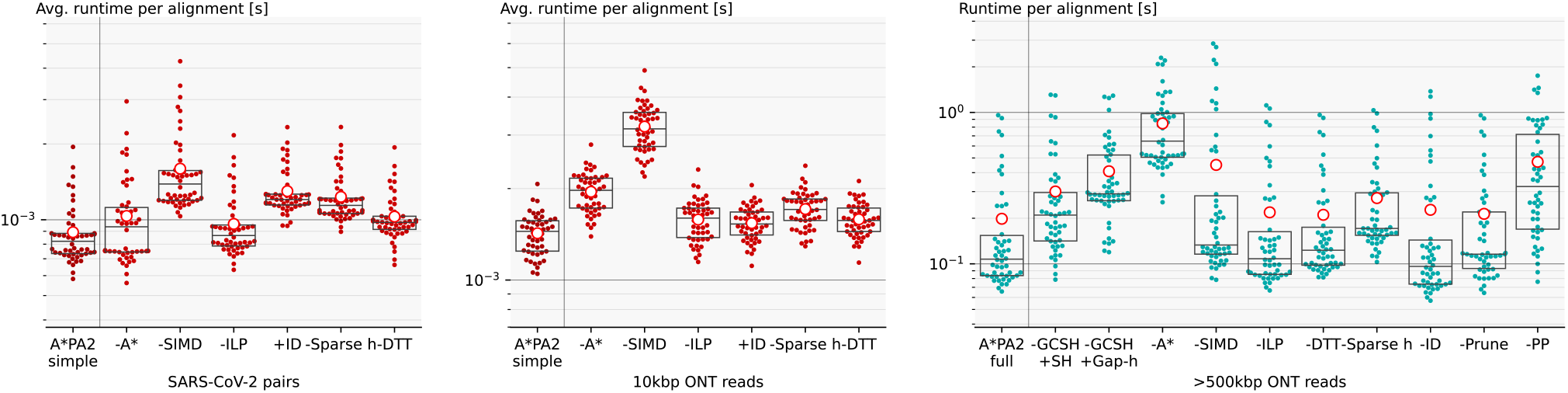
Ablation. Box plots showing how the performance of A*PA2-simple (left, middle) and A*PA2-full (right) changes when removing features. ILP: instruction level parallelims, ID: incremental doubling, Sparse h: spase heuristic evaluation, DTT: diagonal transition traceback, PP: pre-pruning.

**Fig. 13.**
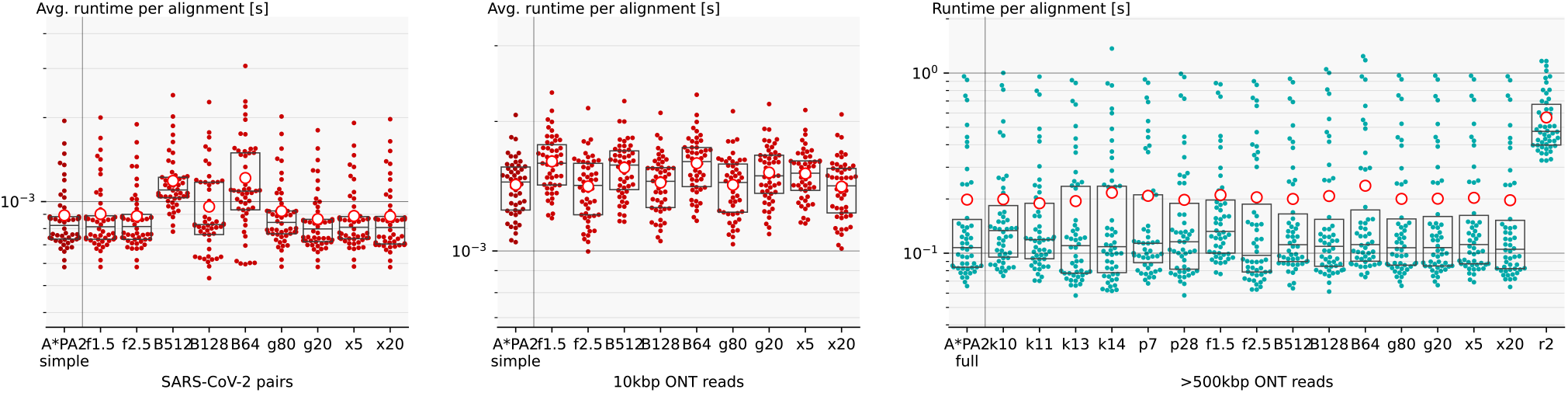
Changing parameters. Runtime of A*PA2-simple (left, middle) and A*PA2-full (right) with one parameter modified. Default parameters are seed length *k* = 12, pre-pruning look-ahead *p* = 14, growth factor *f* = 2, block size *B* = 256, max traceback cost *g* = 40, and dropping diagonals that lag *fd* = 10 behind during traceback. Running time is not very sensitive with regards to most parameters. Of note are using inexact matches (*r* = 2) for the heuristic, which take significantly longer to find, larger seed length *k*, which decreases the strength of the heuristic, and smaller block sizes (*B* = 128 and *B* = 64), which induce more overhead.

**Fig. 14.**
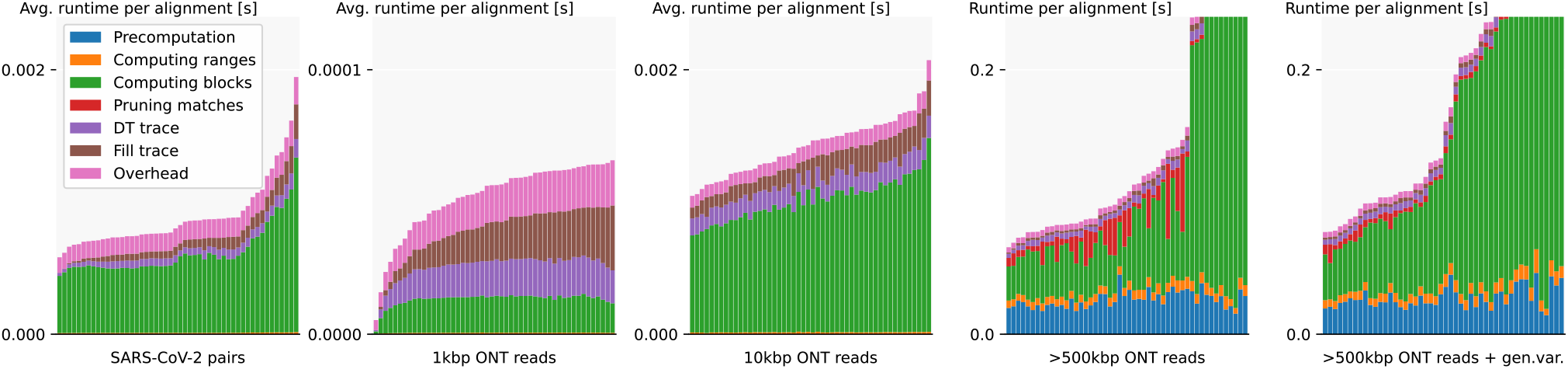
Runtime profile of A*PA2, using A*PA2-simple for short sequences and A*PA2-full for the two rightmost *>*500 kbp datasets. Each column corresponds to a (set of) alignment(s), which are sorted by total runtime. *Overhead* is the part of the runtime not measured in one of the other parts and includes the time to build the profile. For *>*500 kbp long sequences, A*PA2-full spends most of its time computing blocks, followed by the initialization of the heuristic. For very short sequences of 1 kbp, up to half the time is spent on tracing the optimal alignment.

Both methods are usually reported to use fewer than 20 instructions, but exclude the shifting out of the bottom horizontal difference (four instructions) and the initialization of the carry for BITPAL (one operation). We require these additional outputs/inputs since we want to align multiple 64bit lanes below each other, and the horizontal difference in between must be carried through.

### A.2 Sparse heuristic invocation

Here we explain how to compute the fixed range and the range of rows to compute with significantly fewer evaluations of the heuristic than the simpler method of Section 3.8.

#### LEMMA 1.

*lem When h is admissible and f* (*u*) *> t* + 2*D, then f* ^∗^(*u*^*′*^) *> t for all u*^*′*^ *within distance d*(*u, u*^*′*^) ≤ *D from u*.

PROOF. Since adjacent states differ in distance by {−1, 0, +1}, we have *g*(*u*^*′*^) ≥ *g*(*u*) − *d*(*u, u*^*′*^) ≥ *g*(*u*) − *D* and *h*^∗^(*u*^*′*^) ≥ *h*^∗^(*u*) − *d*(*u, u*^*′*^) ≥ *h*^∗^(*u*) − *D*. Now suppose that *f* ^∗^(*u*^*′*^) ≤ *t*. Then *u*^*′*^ is fixed and we have *g*(*u*^*′*^) = *g*^∗^(*u*^*′*^), and since *h* is admissible *h*(*u*) ≤ *h*^∗^(*u*). Thus:

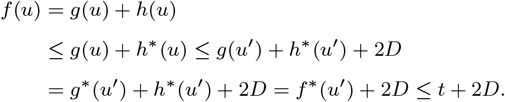

This is in contradiction with *f* (*u*) *> t* + 2*D*, so we must have *f* ^∗^(*u*^*′*^) *> t*, as required.

#### LEMMA 2.

*lem When h is admissible, v is below the diagonal of a computed state u, and f*_*l*_(*v*) = *g*^∗^(*u*) + *c*_gap_(*u, v*) + *h*(*v*) *> t* + 2*D, then f* ^∗^(*v*^*′*^) *> t when v has distance d*(*v, v*^*′*^) ≤ *D from u*.

PROOF. We have *c*gap(*u, v*^*′*^) ≥ *c*gap(*u, v*) − *d*(*v, v*^*′*^) ≥ *c*gap(*u, v*)−*D*, and *h*^∗^(*v*^*′*^) ≥ *h*^∗^(*v*)−*D*. Now suppose that *f* ^∗^(*v*^*′*^) ≤ *t*. Then we know that *g*^∗^(*v*) ≥ *g*^∗^(*u*) + *c*gap(*u, v*), and we still have *h*(*v*) ≤ *h*^∗^(*v*), *g*^∗^(*v*^*′*^) ≥ *g*^∗^(*v*) − *D*, and *h*^∗^(*v*^*′*^) ≥ *h*^∗^(*v*) − *D*. It follows that

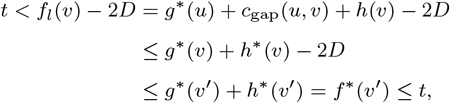

which is a contradiction, so we conclude that *f* ^∗^(*v*^*′*^) *> t*, as required.□

**Step 1’: Sparse fixed range**. To find the first row *jstart* with *f* (⟨*i, jstart*⟩) ≤ *t*, start with *j* = *rstart*, and increment *j* by ⌈(*f* (*v*) − *t*)*/*2⌉ as long as *f* (*v*) *> t*, since none of the intermediate states can lie on a path of length ≤ *t* by Lemma 1. The last row is found in the same way. As seen in Fig. 5b, this sparse variant significantly reduces the number of evaluations of the heuristic in the right-most columns of each block.

**Step 2’: Sparse end of computed range**. Instead of considering one column at a time, we now first make a big jump down and then jump to the right.

1. Start with *v* = ⟨*i*^*′*^, *j*^*′*^⟩ = *u* + ⟨1, *B* + 1⟩ = ⟨*i* + 1, *j*_*end*_ + *B* + 1⟩.
2. If *f*_*l*_(*v*) ≤ *t*, increase *j*^*′*^ (go down) by 8.
3. If *f*_*l*_(*v*) *> t*, increase *i*^*′*^ (go right) by ⌈(*f*_*l*_(*v*) − *t*)*/*2⌉, but do not exceed column *i* + *B*.
4. Repeat from step 2, until *i*^*′*^ = *i* + *B*.
5. While *f*_*l*_(*v*) *> t*, decrease *j*^*′*^ (go up) by ⌈(*f*_*l*_(*v*) − *t*)*/*2⌉, but do not go above the diagonal of *u*.
6. The resulting *v* is again the bottommost state in column *i* + *B* that can potentially have *f* (*t*) ≤ *t*, and its row is the last row that will be computed.

### A.3 Further results

See Figs. 12 to 14 and Tables 1 to 4.

